# Metformin chlorination byproducts in drinking water exhibit marked toxicity to nematode worms, human cells, and mice

**DOI:** 10.1101/2020.07.18.209890

**Authors:** Runshuai Zhang, Yuanzhen He, Luxia Yao, Jie Chen, Shihao Zhu, Xinxin Rao, Peiyuan Tang, Jia You, Guoqiang Hua, Lu Zhang, Feng Ju, Lianfeng Wu

**Affiliations:** School of Life Sciences, Westlake University, 18 Shilongshan Road, Hangzhou 310024, Zhejiang Province, China; Key Laboratory of Growth Regulation and Transformation Research of Zhejiang Province, School of Life Sciences, Westlake University, 18 Shilongshan Road, Hangzhou 310024, Zhejiang Province, China; Institute of Basic Medical Sciences, Westlake Institute for Advanced Study, 18 Shilongshan Road, Hangzhou 310024, Zhejiang Province, China; School of Engineering, Westlake University, 18 Shilongshan Road, Hangzhou 310024, Zhejiang Province, China; Institute of Advanced Technology, Westlake Institute for Advanced Study, 18 Shilongshan Road, Hangzhou 310024, Zhejiang Province, China; Key Laboratory of 3D Micro/Nano Fabrication and Characterization of Zhejiang Province, School of Engineering, Westlake University, 118 Shilongshan Road, Hangzhou 310024, China; Institute of Radiation Medicine and Fudan University Shanghai Cancer Center, Shanghai Medical school, Fudan University, Shanghai 200032, China

## Abstract

Metformin (MET), a worldwide used drug for type 2 diabetes, has been found with the largest amount by weight among all drugs in aquatic environment, including the drinking water sources where chlorination inevitably transforms MET into chlorination byproducts. Although MET has health-promoting properties, whether or how its chlorination byproducts affect health remains largely unknown. Here we reveal that MET chlorination byproducts Y (C_4_H_6_ClN_5_) and C (C_4_H_6_ClN_3_) exhibit marked toxicity, even higher than that of the well-known poisonous arsenic, to live worms and human cells. Moreover, both byproducts are harmful to mice and Y at 250 ng/L destroys the mouse small intestine integrity. Strikingly, we detected MET and byproduct C in worldwide drinking water. Both byproducts are increasingly produced with more MET present during chlorination process. Unprecedentedly, we unveil boiling and activated carbon adsorption as effective solutions that are in urgent demand globally for removing these byproducts from water.

## Introduction

Metformin (MET), as the first-line therapy for type 2 diabetes (T2D), is one of the most prescribed medications in the world. It is anticipated to be consumed in much greater amounts, due to the rapid global increase in T2D prevalence in recent decades and the additionally discovered benefits of MET for cancer prevention (*1*), women’s infertility treatment (*2*) and lifespan extension (*3, 4*). Typically, MET is prescribed with a starting dosage of 500 mg, twice a day for weeks, then increased up to a total of 2,550 mg per day dependent on the tolerance of patients. A latest survey reported that MET was ranked as the 4^th^ most prescribed drug in 2020 and its prescribing rate increased 44.2% from 54.5 million in 2006 to 78.6 million in 2017 in the U.S. (*5*). Meanwhile, China consumes approximately 786 metric tons of MET annually in recent years (*6*). It is noted that MET is not metabolized by human and gets almost 100% excreted unmodified (*7*), resulting in increasing concerns on the potential ecotoxicology of MET and its byproducts after release into the aquatic systems (*8, 9*). Recently, MET present in surface water has been found to be problematic to wild fishes, causing more aggressive behavior (*10*). In fact, MET has already been widely found with the largest amounts among all drugs in waterways and considered as an emerging pollutant (*11*). Although MET could be biotransformed at rates ranging from 41% - 98% in wastewater treatment plants (*12*), it is still widely detected in higher concentrations in surface water (8.70 – 34,000 ng/L) worldwide including many as drinking water sources (Supplementary Table1 and its attached references), necessitating the study of the potential impacts of MET and its derivatives on the safety of drinking water supply and human health.

In the previous MET study (*4*), we incidentally observed an instant reaction of MET with hypochlorite regularly used for *Caenorhabditis elegans* synchronization in the worm field (Supplementary Fig. 2), yielding a yellow product with much higher toxicity than MET itself to nematode worms (data not shown). The reaction was later confirmed and uncovered with two chlorination byproducts of MET: Y (yellow, C4H6ClN5, 159.58 g/mol) and C (colorless, C_4_H_6_ClN_3_, 131.56 g/mol) by Armbruster D *et al (13)*. Chlorine is commonly used for both regional and household drinking water disinfection worldwide owing to its ease of application, low cost, and high efficiency in protecting water from live germ contamination (*14*). We argue that if MET, widely releasing and rapidly accumulating in surface water (*15, 16*), is inevitably transformed into chlorination byproducts of any potential toxicity by chlorination, it will reasonably bring a widespread health threat to all consumers of the water containing such byproducts. MET is thus far the safest drug for diabetes treatment, although it rarely causes toxicity or lactic acidosis when overdose in humans (*17*). In a recent study, the MET byproduct Y was found with genotoxicity in bacteria *Salmonella typhimurium (9)*. However, whether or how MET chlorination byproducts Y and C affect health of animals remains completely unknown. Theoretically, these byproducts are increasingly generated and released into the household taps with more MET present in water sources, but how much they are present in drinking water has not been monitored and paid enough attention yet.

Here, we show that both MET byproducts are even more toxic than arsenic, which is a well-known poisonous compound ever being neglected in drinking water and eventually results in the most serious public health issue historically (*18*), to nematode worms and cultured human cells. Moreover, both byproducts are harmful to mice, and the byproduct Y at likely-achievable doses in tap water remarkably destroys the integrity of the mouse intestinal epithelium in a mechanistic manner likely through the inhibition of intestinal epithelium self-renewal. Continuous disruption of such integrity is known as basis for numerous gut diseases (*19*) and will induce serious and irreversible effects on health if not stopped. Together, we demonstrate the MET chlorination byproducts as a hidden and serious threat to global health and wellbeing. To investigate the present occurrences of MET chlorination byproducts, we examined water samples from household taps of multiple countries and found widespread presences of MET and its chlorination byproduct C in both tap water and drinking water sources. Fortunately, our exploration yields boiling and powered activated carbon adsorption as effective solutions for removing these hazards from drinking water, which are urgently needed for global implementation.

## Results

### MET chlorination byproducts are markedly toxic to live animals and human cells

To demonstrate whether chlorination has transformed MET into toxic byproducts, we optimized the synthesis method (*13*) with low-temperature recrystallization and freeze-drying addition, synthesized and purified the two chlorination byproducts of MET (see methods). Using the firstly purified Y (90.75% of purity) and C (99.51% of purity), we demonstrated that both byproducts at 1 mM (Y = 1.60 × 10^5^ μg/L, C = 1.32 × 10^5^ μg/L) exhibit similar or higher toxicity than MET does at 100 mM (1.29 × 10^7^ μg/L) to *C. elegans* (Fig. 1a), which indicates that chlorination has turned MET into 100 times more toxic compounds to live animals *C. elegans*. Researchers frequently use very high doses and long periods of treatment of MET to induce the effect of interest and seldom consider MET as a toxic reagent. For instance, researchers use over 10 mM (1.29 × 10^6^ μg/L) in cells (*20*), 50 mM (6.45 × 10^6^ μg/L) in *C. elegans* (4, 21) or 150 mg/kg injected intraperitoneally in mice (*22*) generally for 24 or 48 hours in cells and days or weeks in animals (*4, 20, 22*). To define the toxicity level of MET chlorination byproducts, we compared them with arsenic. Strikingly, we found that both MET chlorination byproducts are much more toxic than arsenic when all employed at 2 mM doses, while the byproduct C is 5 times more toxic than arsenic to nematode worms (Fig. 1a).

**Fig. 1.**
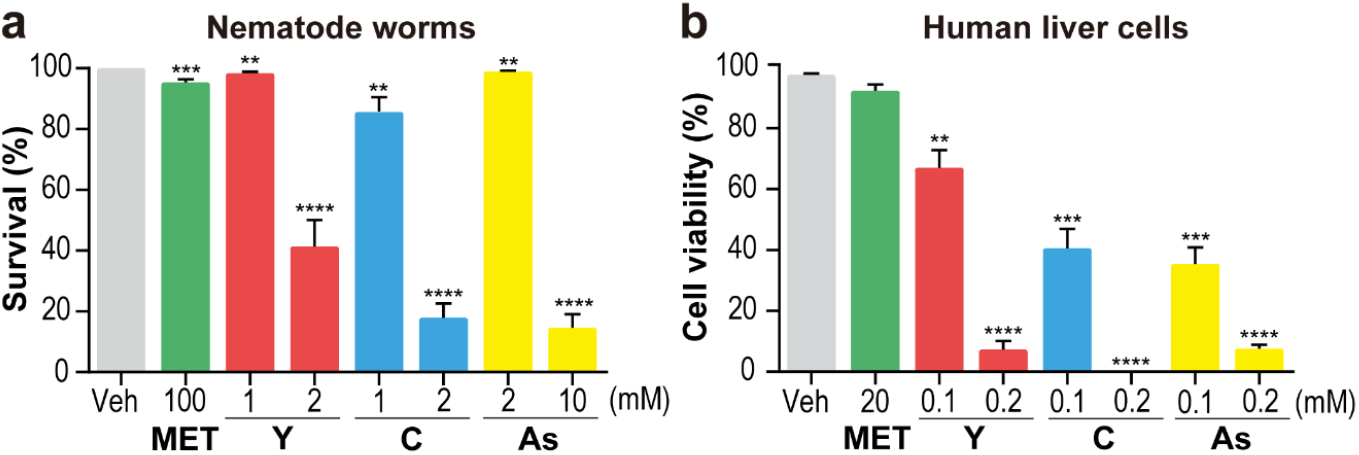
MET chlorination byproducts exhibit marked toxicity to nematode worms and cultured human HepG2 cells, comparable or even higher than that of arsenic (As). **a** Survival rate of larvae L4 worms treated with MET byproducts Y or C (1 mM, Y = 1.60 × 10^5^ μg/L, C = 1.32 × 10^5^ μg/L; and 2 mM, Y = 3.20 × 10^5^ μg/L, C = 2.64 × 10^5^ μg/L) and As (1 mM, 1.3 × 10^5^ μg/L; 2 mM, 2.6 × 10^5^ μg/L) at 50 or 100 times lower concentrations than MET (100 mM, 1.29 × 10^7^ μg/L) for 12 hours. **b** Treatment of MET byproducts Y and C (0.1 mM, Y = 1.60 × 10^4^ μg/L, C = 1.32 × 10^4^ μg/L; and 0.2 mM, Y = 3.20 × 10^4^ μg/L, C = 2.64 × 10^4^ μg/L) for 12 hours resulted in remarkable deaths, which are comparable to the death rates induced by the same doses of As treatment (0.1 mM, 1.3 × 10^4^ μg/L;0.2 mM, 2.6 × 10^4^ μg/L), while MET at 20 mM (2.58 × 10^6^ μg/L) did not cause significant deaths of cultured human liver HepG2 cells. The cell viability was determined by trypan blue staining. The results are presented as the mean SEM of three biological replicates. ** *P* < 0.01, *** *P* < 0.001, **** *P* < 0.0001 by one-way ANOVA.

To assess the cytotoxicity of MET chlorination byproducts, we chose the human liver cell line HepG2, which is widely used in MET studies (*23*) and cytotoxicity tests (*24*). In line with the worm results, we found that MET at 20 mM (2.58 × 10^6^ μg/L) reduced the viability of HepG2 cells slightly, whereas at 0.1 mM (Y = 1.60 × 10^4^ μg/L, C = 1.32 × 10^4^ μg/L) both Y and C induced marked cell death rates and nearly killed all cells at a concentration of 0.2 mM (Y = 3.20 × 10^4^ μg/L, C = 2.64 × 10^4^ μg/L) as measured by trypan blue exclusion assay after 12 hours of treatment (Fig. 1b). To also have an evident sense of the toxicity level of these byproducts in human cells, we compared them with arsenic and found that both MET chlorination byproducts exert similar or even higher toxicity than arsenic to HepG2 cells at the same doses (Fig. 1b). The 50% lethal concentrations (LC50) of Y and C were titrated as 116.5 μM (1.86 × 10^4^ μg/L) and 90.9 μM (1.20 × 10^4^ μg/L), respectively (Supplementary Fig. 3), which are consistently found comparable to the documented arsenic’s LC50 at 100 μM (1.98 × 10^4^ μg/L) in HepG2 cells (*24*).

### MET chlorination byproducts exhibit deleterious effects to mice and their small intestine integrity

To evaluate the toxicity of Y and C at the physiological level, we examined the mouse response to these compounds acutely via a single intraperitoneal injection and chronically through one month of drinking water. In the acute test, we found that all 8 mice injected with 150 mg/kg body weight MET, a regular dose used in MET research (*22*), survived through the 7 day test period and did not show any obvious abnormalities, while 6 of the 8 mice injected with the same dose of C trembled the body, secreted white substances immediately after injection, and died within 2 days (Fig. 2a, Supplementary Fig. 4 and Table 1). However, all 8 mice survived from the 100 mg/kg test of C. In a dramatic contrast, injection of Y at 50 mg/kg led to the deaths of all 8 mice within 2 days and 3 out of 8 deaths at 10 mg/kg 7 days postinjection (Fig. 2a and Supplementary Table 1), which indicates the more potent toxicity of Y than that of C in mice. Moreover, we noticed that the intestines of mice were swollen, dark and bloody after injection of doses higher than 10 mg/kg Y, and swelling was even observed at the low level of 1.25 mg/kg (Fig. 2b and Supplementary Fig. 4), suggesting that Y may attack the mouse small intestines specifically and remarkably.

**Fig. 2.**
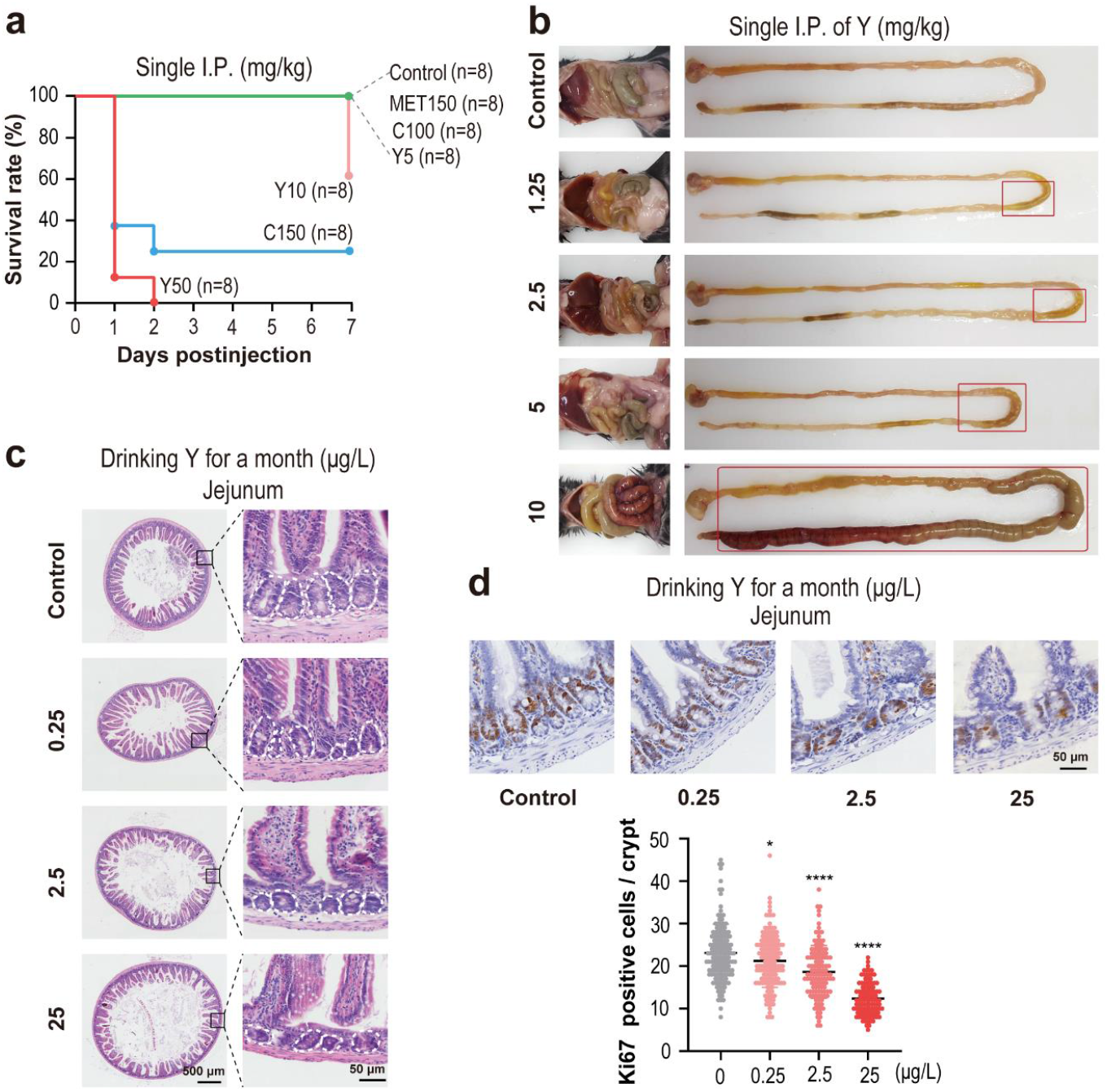
Both MET byproducts Y and C are harmful to mice, and the byproduct Y remarkably destroys the mouse small intestine integrity. **a** Seven-day survival rate of mice after single dose intraperitoneal injection (I.P.) of MET, Y and C at varied doses labeled next to the compound names. For instance, MET150 means the injected dose of MET at 150 mg/kg body weight of the treated mouse. Phosphate-buffered saline (PBS) served as the vehicle for the injected compounds and the injection control. The number of mice (n) injected in each group is provided in parentheses. Detailed observations of the mice after injection and more acute toxicity groups are attached in Supplementary Fig. 4 and Table 3. **b** Single-dose injection of Y at doses of 1.25 mg/kg and higher led to severe mouse small intestine abnormalities. Surviving mice were examined for tissue or organ alterations by different doses of Y on day 7 after injection. Small intestines of representative treated groups were isolated to show the abnormalities. In the chronic response test for **(c)** and **(d)**, all mice that survived from drinking water with byproduct Y (0.25, 2.5, and 25 μg/L) or without for one month were dissected for evaluation by hematoxylin-eosin (H&E) staining and Ki67 immunohistochemistry (IHC) staining at the end of exposure. Images are representative of 8 animals for each group. **c** A month of exposure to Y at 2.5 μg/L and 25 μg/L caused marked morphological changes in the small intestine crypts, circled in white dashed lines. **d** Byproduct Y inhibits the proliferation of crypt cells in the jejunum in a dose-dependent manner. Quantification of the Ki67-positive cells in each crypt was conducted and is shown next to the IHC images. Bars represent the SEM (n=100-150 crypts from 8 mice). * *P* < 0.05, *** *P* < 0.001 by one-way ANOVA.

Chronic response tests were performed to determine whether low doses of Y and C could cause adverse effects in mice through daily exposure in drinking water for one month. After one month of drinking the water containing 2.5 mg/L C, the mice did not show any obvious abnormal behaviors (data not shown) or intestinal morphology changes as determined by histological staining (Supplementary Fig. 5). However, consistent with the acute test results, we observed that the byproduct Y caused morphological changes in the crypts at the jejunum of the small intestine, where the majority of nutrients are absorbed (*25*), after one month of treatment in drinking water at a dose of 0.25 μg/L (Fig. 2c). As documented, most cells at the base of small intestine crypts are stem cells that normally turn over every 3 to 4 days for cell renewal of the epithelium (*26, 27*). To determine whether Y induces intestinal epithelial layer changes by affecting cell proliferation of crypt cells, we conducted an immunostaining assay against the protein Ki67, a marker of cell proliferation (*28*). According to Ki67 staining, Y inhibited the proliferation of cells at the base of the small intestine crypt in a dose-dependent manner (Fig. 2d), which implicates that Y impedes self-renewal of the intestinal epithelium by blocking cell proliferation. The continuous disruption of the intestinal epithelium by the byproduct Y is likely to induce serious or irreversible threats to health and wellbeing.

### Current contamination and projected future accumulation of MET and its chlorination byproducts in the global drinking water

Given that the toxicities of MET byproducts to nematode worms and cells are comparable or higher than arsenic and the byproduct Y impairs the integrity of the mouse small intestine when administered in drinking water, we should not experience any epidemiological disaster like that was caused by arsenic contamination in drinking water, which has been recognized as a historical mistake by ignoring arsenic in drinking water quality control (*29*). To ascertain the current presences of MET and its chlorination byproducts in drinking water, we employed ultra-high performance liquid chromatography-triple quadruple tandem mass spectrometry (UPLC-MS/MS) to examine water samples from household taps worldwide, especially where surface water is used for a drinking water source. Markedly, we found that the current concentration of MET ranges from 5.12 to 1,203.50 ng/L, and byproduct C ranges from 0.32 to 9.72 ng/L in urban drinking water from multiple countries, including China and the U.S. (Fig. 3a and Supplementary Table 2). The detection limit of byproduct Y is relatively high by UPLC-MS/MS (90.75 ng/L) possibly because Y is labile under the high temperature of the ion source of the mass spectrometer (Supplementary Fig. 6) and as reported by Armbruster D *et al (13)*.

**Fig. 3.**
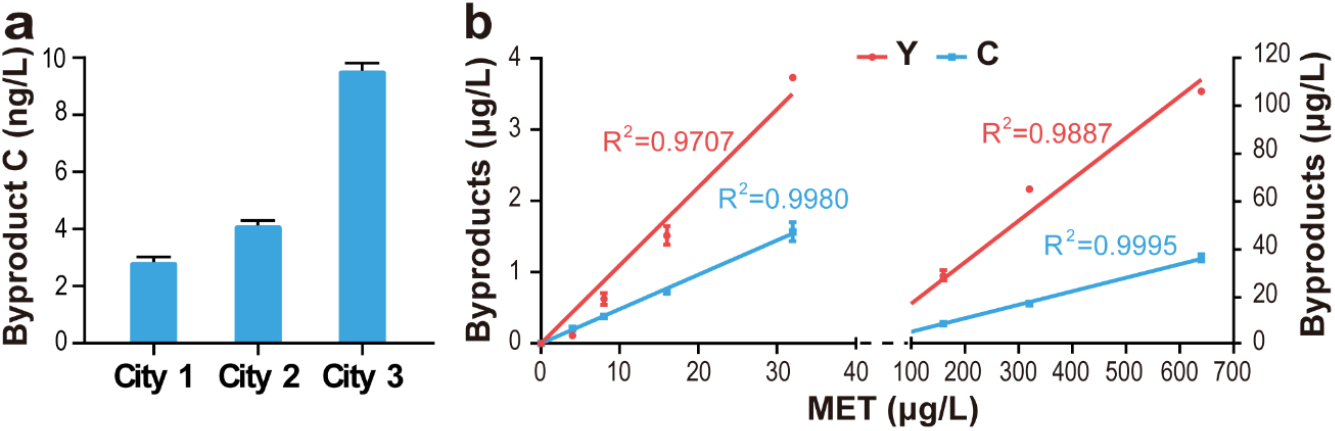
Current and predicted future presence of MET chlorination byproducts in drinking water. **a** MET byproduct C is widely detected in household tap water. The bar graph represents results of three biological samples from the same place of the top three C-rich cities examined. The error bar represents the SEM. **b** A simulation experiment was carried out to depict the theoretical production of MET chlorination byproducts with a fixed concentration of chlorine (0.3%) regularly used in water treatment and an increasing addition of MET. The byproduct Y is colored in red and C is colored in light blue.

Although the current concentration of Y was not determined in tap water samples in this study, possibly due to its detection limit on the UPLC-MS/MS system, C has been widely detected in tap water from multiple countries and even found in a lake water from China at a concentration of 3.46 ng/L, suggesting that the MET byproduct C has entered into the water cycling from household taps to surface water resources (Supplementary Table 2). Accumulating evidence shows that MET typically exceeds 1 μg/L in surface water worldwide (Supplementary Fig. 1) and is particularly high in some surface waters in the U.S. (*16*), China (*30*) and Canada (*31*) as 34 μg/L, 20 μg/L and 10 μg/L, respectively. Unfortunately, it is noteworthy that many countries, including the U.S. and China, take surface water for drinking water generation without bank filtration or any other underground passage process, which are thought to be efficient ways to reduce MET from surface water (*32*). To predict the presences of these chlorination byproducts in tap water with increasing MET levels, we carried out simulated disinfection experiments and found steady growth of Y and C with increased addition of MET during chlorination process (Fig. 3b). Thus, theoretically, the byproduct Y in tap water is most likely to achieve health-threatening doses over time unless the MET disposal or clearances of its chlorination byproducts are well controlled.

### Solutions for removing MET chlorination byproducts from water

The reason why we do not see the havoc causality from MET chlorination byproducts in drinking water to potential health risks yet may lie in our means to use tap water, such as drinking the water after boiling, which is regularly employed in many countries, including China. We surprisingly found that boiling makes the byproduct Y vanish and C significantly reduced from water (Fig. 4) and turns the byproduct Y safe to mice even at the highly lethal dose (50 mg/kg Y, I.P., Supplementary Fig. 7), although we do not know what boiling transforms them into. It implies that boiling could be one option to avoid the potential risks from drinking the water containing MET chlorination byproducts, especially for the harmful Y. To prevent MET chlorination byproducts in tap water from reaching the harmful levels, we exploited the possibility that activated carbon adsorption, which is commonly used in water treatment, might help to remove these compounds from water. Activated carbon adsorption is a well-established approach in removing disinfection byproducts generated in water treatment due to its well-known effectiveness and the ease of activated carbon regeneration (*33, 34*). We found that powdered activated carbon adsorbs byproducts Y and C more effectively than the granular form does (Fig. 4), the results of which can be leveraged to guide water treatment routes and develop additional processes for completely removing Y and C from drinking water.

**Fig. 4.**
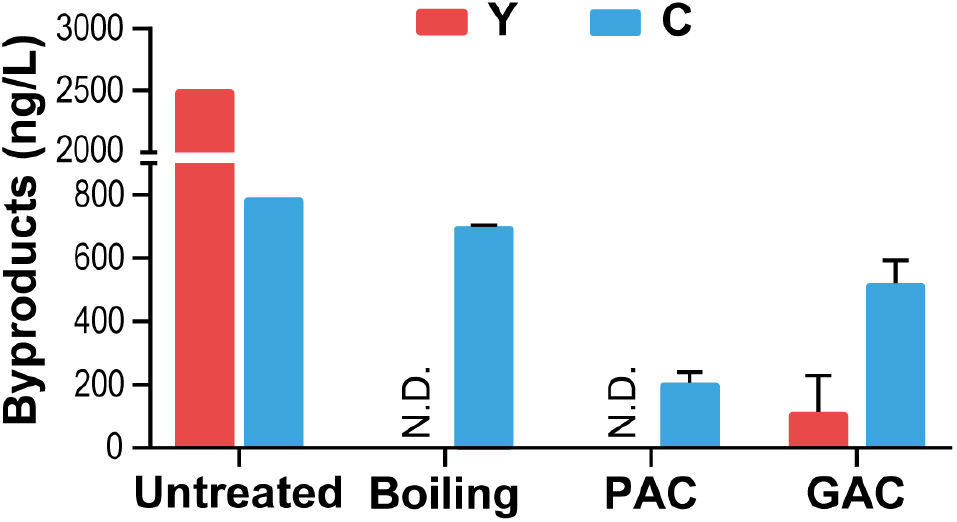
MET chlorination byproducts can be removed from water by boiling and activated carbon adsorption. Boiling makes byproduct Y vanish, while activated carbon, including powdered activated carbon (PAC) and granular activated carbon (GAC), can effectively remove both MET byproducts Y and C from water. MET byproducts Y and C in water were analyzed by UPLC-MS/MS prior to and after treatment, including boiling and adsorption through PAC or GAC. N.D. stands for not detected.

## Discussion

MET has been widely recognized as a panacea to promote health, while our findings here reveal that hypochlorite turns this panacea drug into toxic players (byproducts Y and C) with demonstrated harm to the health of life over time (Supplementary Fig. 8). Although the current doses of MET present in drinking water may not directly cause safety concerns to humans, the potential threats of its chlorination byproducts cannot be neglected. Because they were found to be toxic to live animals *C. elegans* and mice at doses generally safe for MET administration, and have been traced in household tap water consumed daily by billions of people globally, including young and sick people. Concluding from the historical lessons of arsenic contamination in drinking water, Smith AH *et al.* highlighted that “delaying action in an attempt to be thorough in research and long-term planning can be a mistake” (*29, 35*). Our current study necessitates the emergent need to globally pay attention to the MET byproducts widely present in chlorinated drinking water.

Previous lessons have taught us that drinking water containing 50 μg/L arsenic would cause 1 in 100 people to die from arsenic-related cancers, including lung, bladder, and skin cancers (*35*). It is noteworthy that both MET chlorination byproducts Y and C are highly nitrogenous substances, which is a typical trait of bladder carcinogens identified to date(*36*). Whether these previously neglected MET byproducts are potential bladder carcinogens, or carcinogen precursors needs to be explored. Especially, whether or how its byproduct C affects the health of mice and wild animals needs to be further explored, as it is stable and has been widely detected in tap and surface water.

To our surprise, MET byproduct Y induced significant toxicity to mice in both the acute test and the chronic treatment, especially attacking the mouse gut. The reason why compound Y targets the gut remains elusive, but collective evidence has proposed the gut as a major action site of MET, increasing glucose uptake and lactate production within the intestine(*37*). More importantly, MET has been found to be mostly absorbed in the small intestine and highly accumulated in the jejunum peaks (*38*). It is plausible that the compound Y is also adsorbed in the small intestine and thus acts directly in the gut.

Although boiling and activated carbon adsorption can be used as feasible solutions to remove the chlorinated byproducts of MET from water, immediate measures should be taken to stop the growing health threats to humans and wildlife from daily exposure of these byproducts in drinking water. Strategies to remove MET from the environment were intensively explored (*32, 39, 40*), but very few effective ones have been suggested, including bank filtration or underground passage (*32*) and biodegradation by certain species of bacteria (*40*). Alternatively, additional, better drug options to replace MET or reduce MET consumption are also in emerging demand.

## Methods

### Synthesis and purification of MET chlorination byproducts Y and C

MET chlorination byproducts were prepared according to a protocol modified from the previous study by Armbruster D *et al (13)*. Briefly, a series of steps were carried out including chlorination reaction of MET (sigma, PHR1084), extraction, rotary evaporation concentration, column chromatography purification and recrystallization. In terms of the potential heat-labile property of the byproduct Y, low-temperature recrystallization (4 °C in the dark) was additionally applied to get the compounds Y and C of maximal purity, followed by separation of the solid-liquid phase and further removal of water through freeze-drying (−60°C, 12 hours). The purities of synthesized byproducts Y (bright orange powder) and C (colorless powder) were determined by nuclear magnetic resonance (NMR), as 90.75% and 99.51% respectively. The synthesized Y and C were then used as standards for mass spectrometry analysis of their concentrations in global surface water and drinking water samples, and for the treatment of nematode worms, cultured cells, and mice. Consistent with the previous observation that Y is partly converted to C during preparation (*13*), a certain amount of C signal (4.48%) was detected in the purity analysis of Y, which is also reflected in the same results of mass spectrometer analysis (Supplementary Fig. 6) and explains why the purity of Y is lower (90.75% vs. 99.51%).

### Cell culture and cell viability test

HepG2 cells were cultured in Dulbecco’s Modified Eagle’s Medium-High Glucose (DMEM) medium containing L-glutamine (4 mM; Sigma) and 1 mM sodium pyruvate solution (100 mM, Sigma), 10% fetal bovine serum (FBS) and 1% penicillin-streptomycin. For the cytotoxicity test, the cells were seeded in 24 wells plate at 5×10^4^ per well overnight before drug treatment. The cell viability was determined by trypan blue staining after the 12-hour treatment of the tested compounds at indicated dose.

### Worm culture

The wild-type *C. elegans* strain N2 was maintained at 20 °C on nematode growth media (NGM) plates that were seeded with *Escherichia coli* (*E. coli*) OP50 as a food source. Toxicity tests were conducted with L4 larvae in M9 buffer with different concentrations of MET (sigma, PHR1084), sodium arsenite (sigma, S7400), and the byproduct Y or C.

### Mice

Female C57BL/6 mice at 8-10 weeks old were used for acute and chronic toxicity tests. For the acute toxicity test, the compounds were diluted in PBS and administered intraperitoneally. Animals were monitored and recorded on survival and behavior changes for 7 days. For chronic toxicity test, compounds were delivered through drinking water for one month. Drinking water containing byproduct Y or C is prepared freshly every day. Mice were maintained and handled in accordance with institutional guidelines, and all animal procedures were approved by the Institutional Animal Care and Use Committee of Westlake University.

### Tissue preparation for immunohistochemistry (IHC)

Dissected intestinal tissues were thoroughly rinsed twice before being fixed in 10% neutral-buffered formalin (Leagene, DF0111) for 24 hours and embedded in paraffin blocks. Paraffin sections (5 μm) were stained with hematoxylin/eosin (H&E; Merck, Darmstadt, Germany). For immunohistochemistry (IHC), sections were boiled in 0.1 M citrate buffer (PH 6.0) 30 min for antigen retrieval. The sections post boiling were incubated with Ki67 antibody (Abcam, ab16667; dilution 1:500) overnight at 4 °C in a humidified chamber followed by detection employing UltraSensitiveTM SP (Mouse/Rabbit) IHC Kit (Maxim, KIT-9720) and DAB kit (Maxim, DAB-1031) according to manufacturer’s instructions. All stained sections were scanned using TissueFAXS microscope (TissueFAX plus; TissueGnostics, Vienna, Austria).

### Water sample collection

The tap water sample or its sources from various lakes or rivers were collected from different cities of China, the U.S., Japan, Korea, and Philippines from June to September in 2019. All water samples were collected freshly and refrigerated at −20°C for later transportation and storage. Samples were filtered with 0.22 μm mixed cellulose (MCE) membrane prior to analysis by UPLC-MS/MS.

### UPLC-MS/MS trace analysis

UPLC-MS/MS (RxionLC™ SCIEX 6500+, QTRAP, SCIEX, U.S.) in the positive electrospray ionization (ESI+) mode was used to measure the trace amounts of MET and its byproducts Y and C. MET separation was achieved on an BEH Amide analytical column (50 mm×2.1 mm, 1.7 μm; Waters, U.S.) using a mixture of 0.1% formic acid aqueous solution containing 5 mM ammonium acetate (eluent A) and acetonitrile (eluent B: ACN). Gradient elution was set up as: 0 min (95% ACN); hold: 1 min (95% ACN); ramp: 4 min (40% ACN); hold: 5 min (40% ACN); ramp: 5.5 min (95% ACN); stop: 7 min (95% ACN). Y and C in the collected samples were separated by use of a reversed phase column (ZORBAX Eclipse plus C18, 50 mm×2.1 mm, 1.8 μm; Agilent, U.S.) with a gradient of 95 % eluent A (ultrapure water with 0.1% formic acid), hold for 1 min, rise to 90% eluent B (Methanol plus 0.1% formic acid) in 2 min, hold for 1 min, rise to 95% eluent A in 0.5 min and hold for 2 min (total time for 7 min). The UPLC was operated at a flow rate of 0.4 mL/min, with injection volume of 20 μL and column temperature at 40 °C. The optimized mass spectrum conditions were run as follows: source temperature at 500 °C; IonSpray voltage at 2,000 V; curtain gas at 35 psi; Ion source gas 1 and 2 at 55 psi and 50 psi, respectively. The MET was quantified based on the external calibration using the MET (Sigma, PHR1084) as standard and its recovery ranged from 98% to 126% with quantification based on external calibration and recovery. The limits of detection and quantification (LOD and LOQ) for MET were 2 ng/L and 5 ng/L, respectively. Our synthesized byproducts Y and C were used as standards to analyze water samples collected. The recovery of the byproduct C from the water samples was 86.87% (mean), and its LOD and LOQ were determined as 0.2 ng/L and 0.5 ng/L, respectively. The quantification of Y was achieved by multiple reaction monitoring (MRM) chromatogram of C (132.20 to 71.00; the retention time of C is at 2.47 min) as it was also present at the retention time of Y (2.70 min) after passing through the ion source (Supplementary Fig. 6). The signal decomposition from Y to C appeared to be linearly correlated well. Thus, the LOD and LOQ of Y were determined as 72.60 ng/L and 90.75 ng/L respectively (Supplementary Fig. 6).

### Disinfection assay with water containing MET

To simulate the formation of MET byproducts Y and C during chlorination, NaOCl (Aladdin, Shanghai, China, Cat. S101636) was used to react with MET in ultrapure water (Milli-Q system) in triplicates. In brief, 1.5 mL NaOCl solution with 100 mg/L active chlorine was added into 50 mL glass tube containing 48.5 mL of metformin solution in the range of 4 μg/L (currently detected level in the drinking water source of this study) to 640 μg/L. The chlorine concentration was within the limit of Standards for Drinking Water Quality in China (GB 5749-2006, 2006). The reaction was carried out in dark and on stirring plate at low speed for 30 min to ensure the enough reaction time. At the end of reaction (2 h), the reaction mixtures were quenched with 1 μL, 10 g/L sodium thiosulfate (Na2S2O3) followed by transferring into 1.5 mL brown sample vials. Collected samples were stored in the dark at −20°C and analyzed within 3 days.

### Boiling test

100 mL Y or C solutions, at a concentration of 2,509.50 ng/L and 790.41 ng/L respectively, in 250 mL beakers were heated by an electric water heater. Beakers were sealed with foil to minimize evaporation loss of solutions. The samples were immediately collected into 1.5 mL brown sample vials after boiling, and frozen for storage at −20°C before analysis. Triplicated experiments were conducted, and concentrations of Y and C before and after boiling were determined via UPLC/MS/MS.

### Activated carbon adsorption

Two forms of activated carbon with different particle sizes (200 mesh and 1-2 mm; Huanyu Carbon Industry CO., LTD. China) were evaluated as the potential sorbents of MET chlorination byproducts according to the manufacturer’s instructions. Both powdered and granular activated carbon were dried in oven at 80 °C for 5 hours to remove water. In each test, 100 mg activated carbon was added to 30 mL Y solutions (2,509.50 ng/L) or C solutions (790.41 ng/L) for adsorption followed by 24-hour equilibration in a constant temperature shaker (25 °C). At the end of equilibration, activated carbon was removed from Y or C solutions with 0.22 μm MCE membrane filters. The concentrations of compound Y or C remaining in solutions were subsequently determined by UPLC-MS/MS. All reactions were carried out in brown glass vials.

### Statistical analysis

For cell viability test, data were obtained from 3 biological replicates. For Ki67 staining assay, at least 5 images for each condition were quantified and analyzed. Experimental differences were analyzed by one-way ANOVA in Prism 7.0 d. Values represent mean ± SEM. P value ≤ 0.05 is considered of significance.

### Data availability

All data supporting the findings of this study are available within the paper and its supplementary information file.

## Acknowledgments

We thank Jinheng Pan, Shan Feng, Nanjia Zhou, Hang Shi and Xiaohuo Shi for the facility support and discussions. We are grateful to all people volunteered for assistance on water sampling. We thank Alexander A. Soukas, Dangsheng Li, Ling Li, Yigong Shi, Tian Xu, and Li Deng for critical discussions. This work was supported by institutional funds from the Westlake University / Westlake Institute for Advanced Study (L.W. and F.J.), by the Young Scientists Fund of the National Natural Science Foundation of China, grant number 51908467 (F.J.), by the National Natural Science Foundation of China, grant number 31670858 (H.G.).

## Author Contributions

R. Z., Y. H., L. Y., F. J. and L. W. designed the experiments. R. Z., L. Y., J. C., S. Z., X. R., P. T. and Y. J. performed the cell and mouse experiments. Y. H. and L. Z. synthesized the compounds and performed water sample analysis. J. Y. and G. H. mentored the mouse experiments and analyzed the results. R. Z., Y. H., L. Y., F. J. and L. W. wrote the manuscript. F. J. and L.W. supervised the project.

---

**Supplementary Figure 1.**
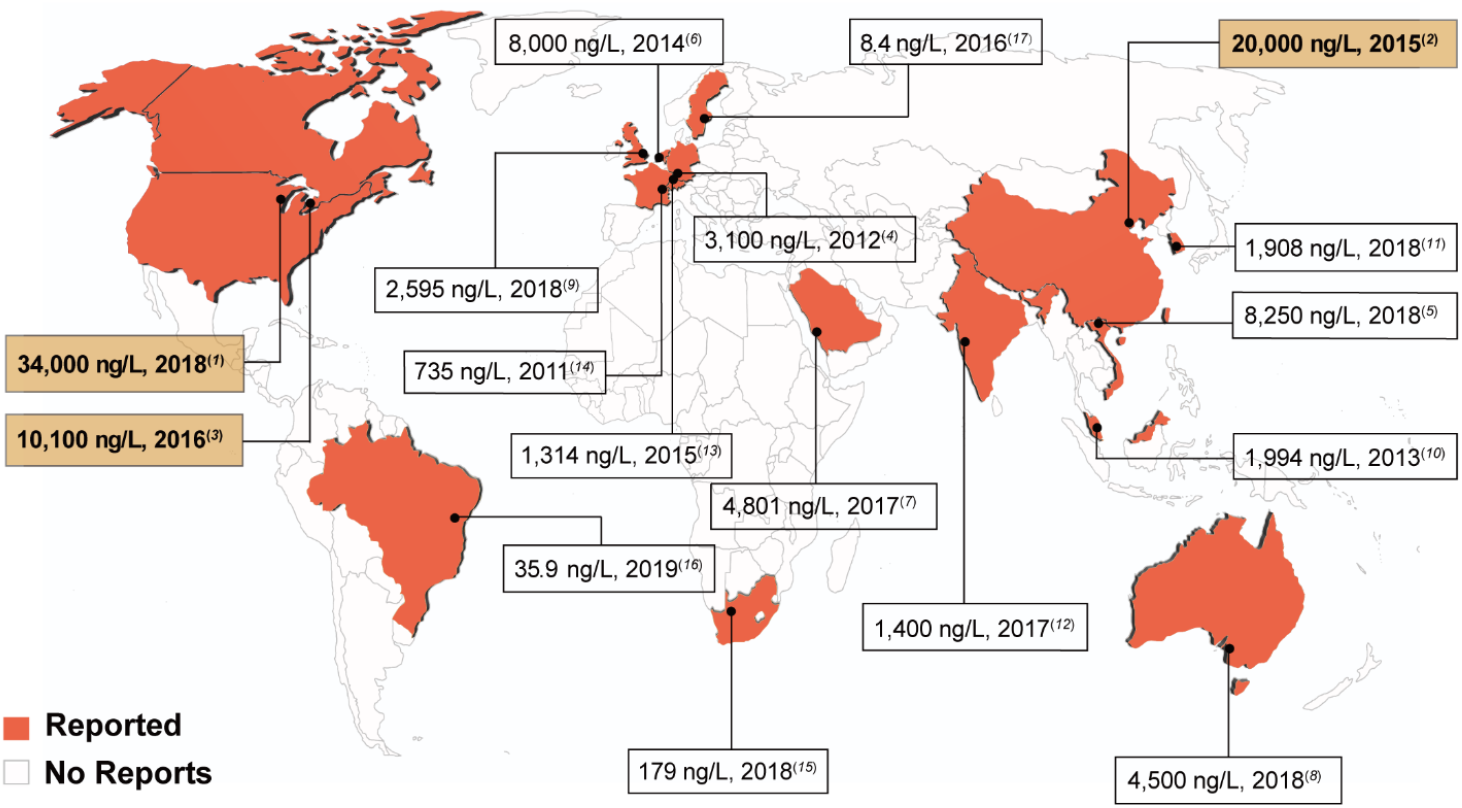
Global view of the occurrence and concentration of MET currently reported in surface water. The black spots on the map represent the reported sampling sites in the corresponding countries. The reported concentration and year are provided and labeled with related references in each box, where the three spots are highlighted in orange with the highest concentration of MET reported in surface water.

**Supplementary Figure 2.**
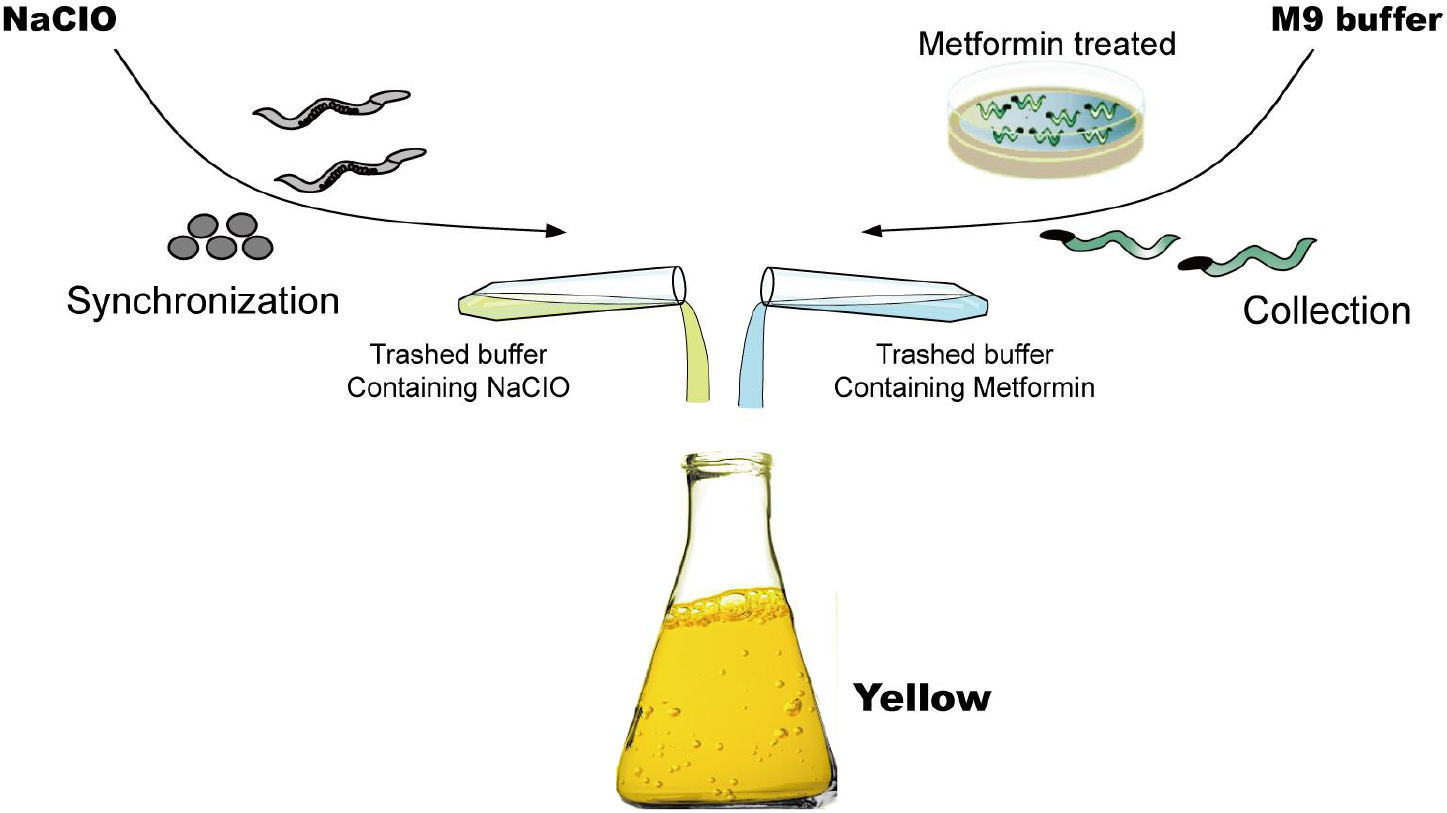
An incidental observation leading to the first knowledge of MET chlorination byproducts with potential toxicity of to the nematode *C. elegans.* An instant reaction occurs in mixed trash buffer from worm synchronization and washes of MET treated worms.

**Supplementary Figure 3.**
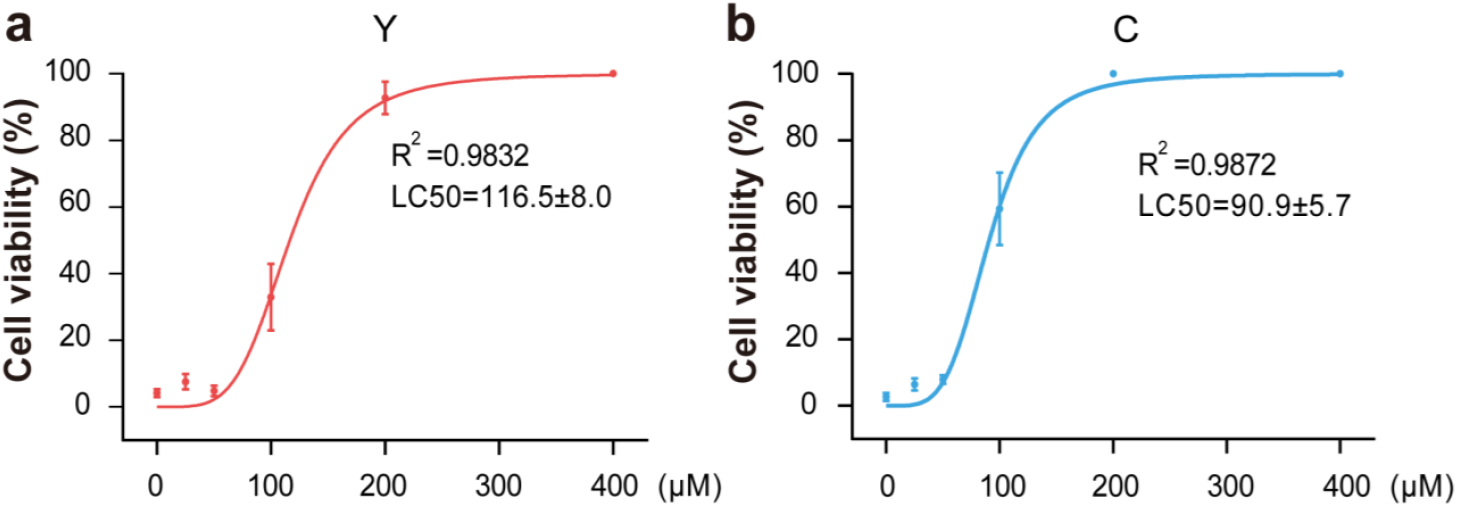
Correlation curves of cytotoxicity and dose of byproducts Y and C were generated from trypan blue staining assays in HepG2 cells. The 50% lethal concentrations (LC50) of (A) Y and (B) C that are lethal to 50% of tested cells were determined from the assay as 116.6 ± 8.0 μM (1.86 × 10^4^ μg/L) and 90.9 ± 5.7 μM (1.20 × 10^4^ μg/L), respectively. Data are presented as the mean ± SEM for 3 biological replicates.

**Supplementary Figure 4.**
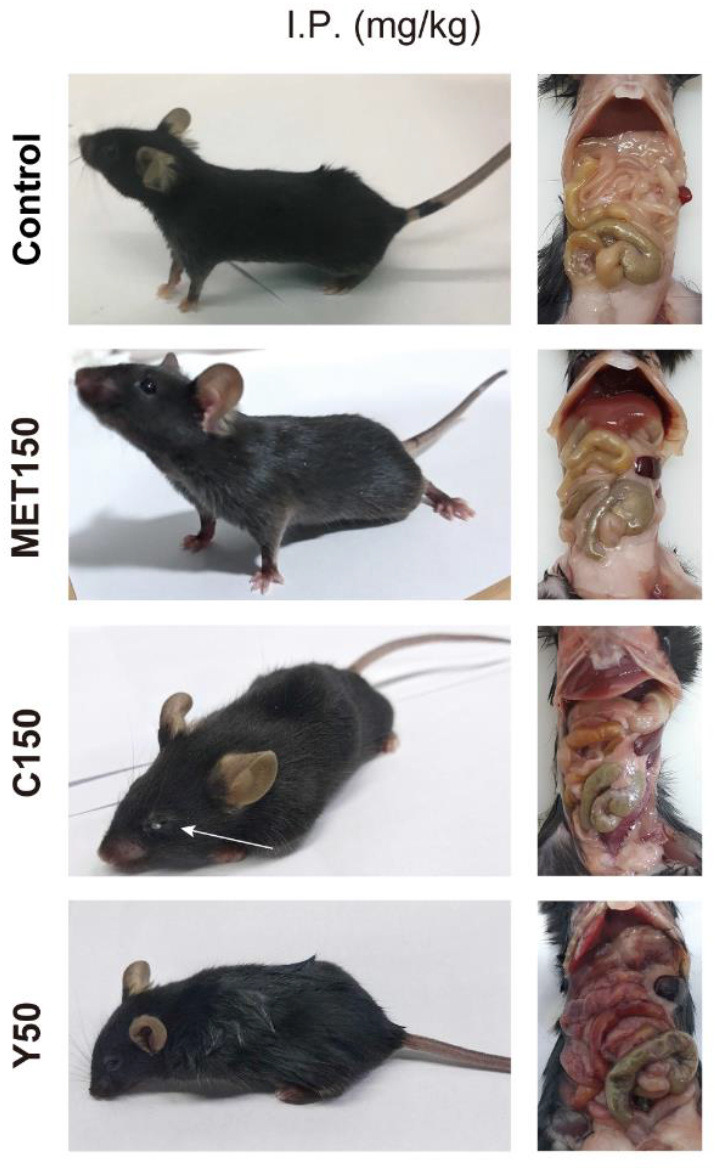
Acute responses of mice to the tested compounds in minutes or hours (pictures on left) and after 7 days (dissection images on the right) after intraperitoneal injection. PBS was injected as a control. MET was given at a dose of 150 mg/kg body weight of mice, byproduct C at a 150 mg/kg dose and byproduct Y at a 50 mg/kg dose. The white arrow is used to draw attention to secreted white substances from the eyes of the mice 10 min after injection of byproduct C, which disappeared within 2 hours. All 8 animals injected with Y at 50 mg/kg died within 2 days, the representative image of which shown on the right was taken on the day of death.

**Supplementary Figure 5.**
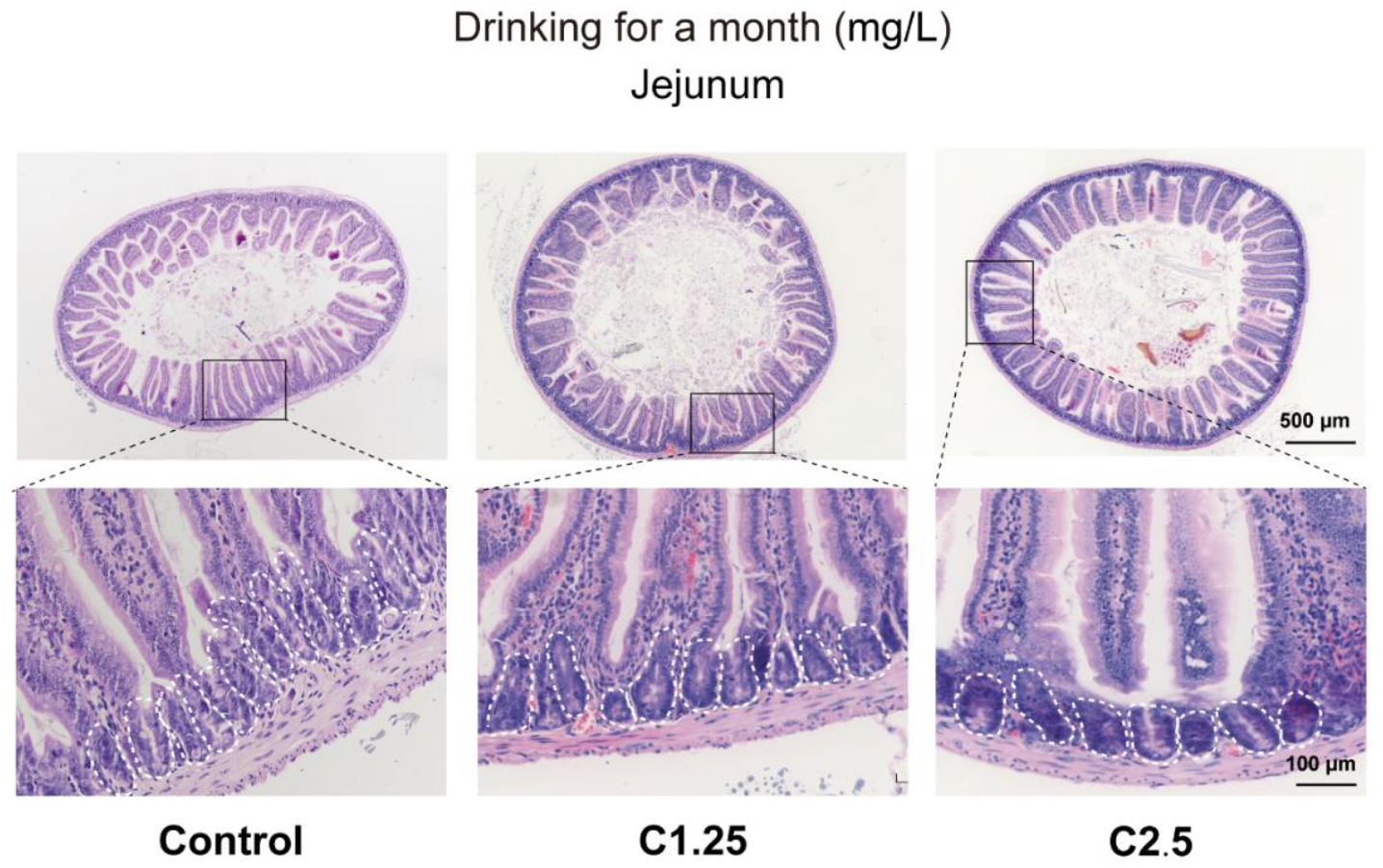
No obvious morphological changes in the small intestine were observed in mice exposed to byproduct C through drinking water for one month. Images from the H&E staining assay are representative of 8 animals from each group. Crypts are circled with white dotted lines. Water served as a control, and C was administered by feeding water containing C at the level of 1.25 mg/L or 2.5 mg/L.

**Supplementary Figure 6.**
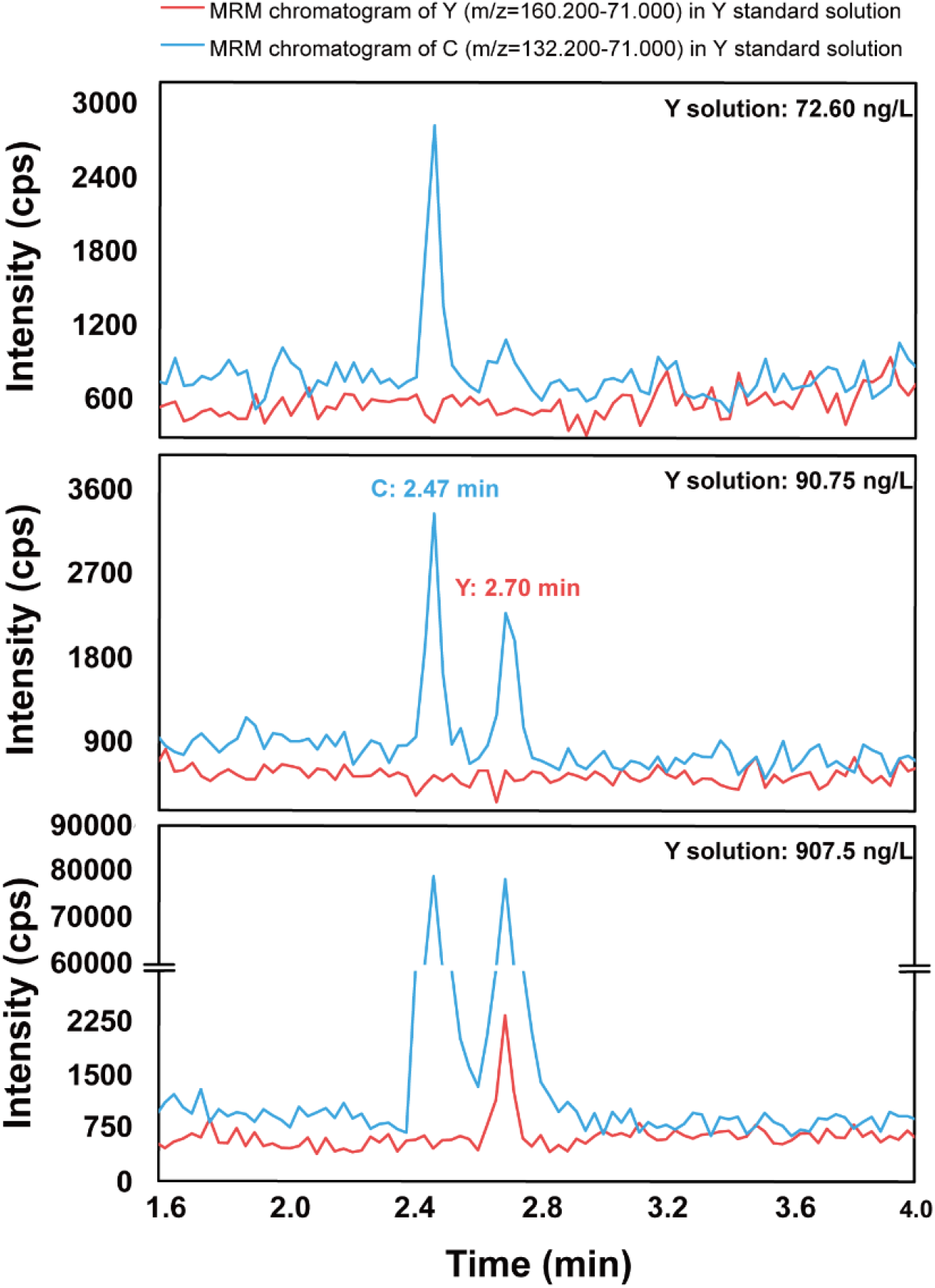
Determination of the detection limit of byproduct Y by UPLC-MS/MS. The X axis represents the time of compound flowing through, while the Y axis represents the intensity of the flowing compound. The retention times of compounds Y and C are 2.70 min and 2.47 min, respectively. As shown in the plot, byproduct Y is transformed into C after injection when going through the ion source of the mass spectrometer as the “C peak” also appears at the retention time of Y in multiple reaction monitoring (MRM) of C in the UPLC-MS/MS result of Y standard test. In addition, the peak at 2.70 min (signal from Y to C) disappears when Y is injected at a low concentration (< 72.60 ng/L) from the upper plot. The signal of Y to C starts to show and be quantifiable by mass spectrometry at a concentration of 90.75 ng/L and becomes significant at 907.50 ng/L, as indicated by the middle and bottom plots.

**Supplementary Figure 7.**
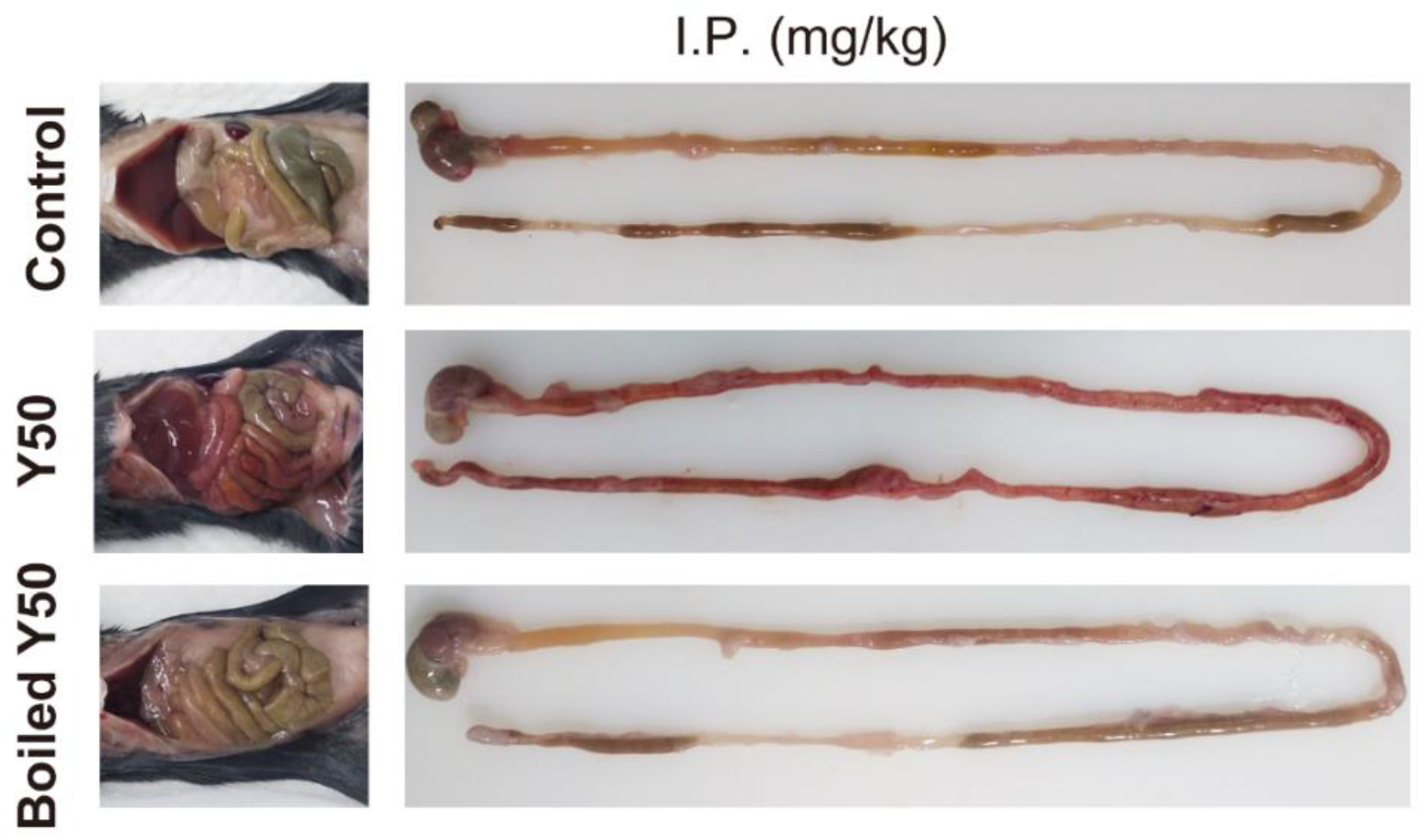
MET byproduct Y after boiling at the extremely toxic dose of 50 mg/kg by I.P. does not induce any obvious effects in mice. PBS after boiling served as injection control (5 animals). MET byproduct Y, boiled, or unboiled, was injected at the dose of 50 mg/kg body weight of mice (8 animals each group). All 8 animals injected with unboiled Y died within 2 days, the representative image of which was taken on the day of death, while animals in the control and boiled groups were sacrificed and dissected for analysis at the end of 7 days of evaluation.

**Supplementary Figure 8.**
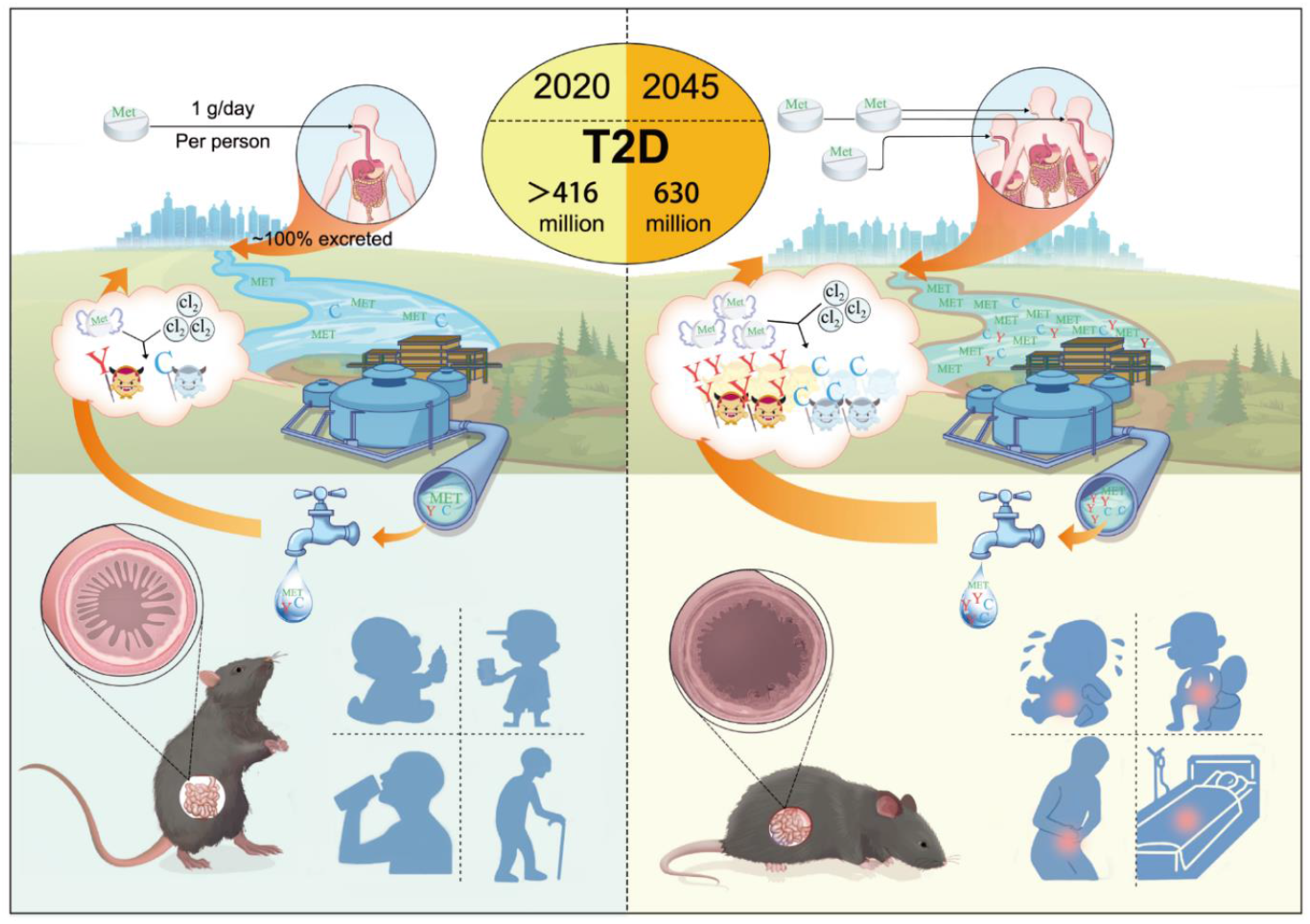
Projection model of the fast-growing threat from daily exposures of MET chlorination byproducts in drinking water. Regularly, MET is prescribed to individuals with T2D at approximately 1 g/day/person, and almost 100% is excreted from the body. Surface water containing MET, as a main drinking water source worldwide, is most often disinfected with chlorine or hypochlorite prior to household usage. The reaction occurs instantly, as displayed in the cloud chart, and MET is turned into Y and C in a MET dose-dependent manner during the chlorination process. The T2D number provided in the middle pie graph is calculated according to the 9th edition of the International Diabetes Federation, reporting 463 million people currently with diabetes worldwide and presumably up to 700 million in 2045, approximately 90% of whom have T2D^18^. With daily exposure to growing doses of MET byproducts in the future, mice, and humans of various ages and in various conditions are depicted to highlight potential risks of attack on the gut.

**Supplementary Table 1.**
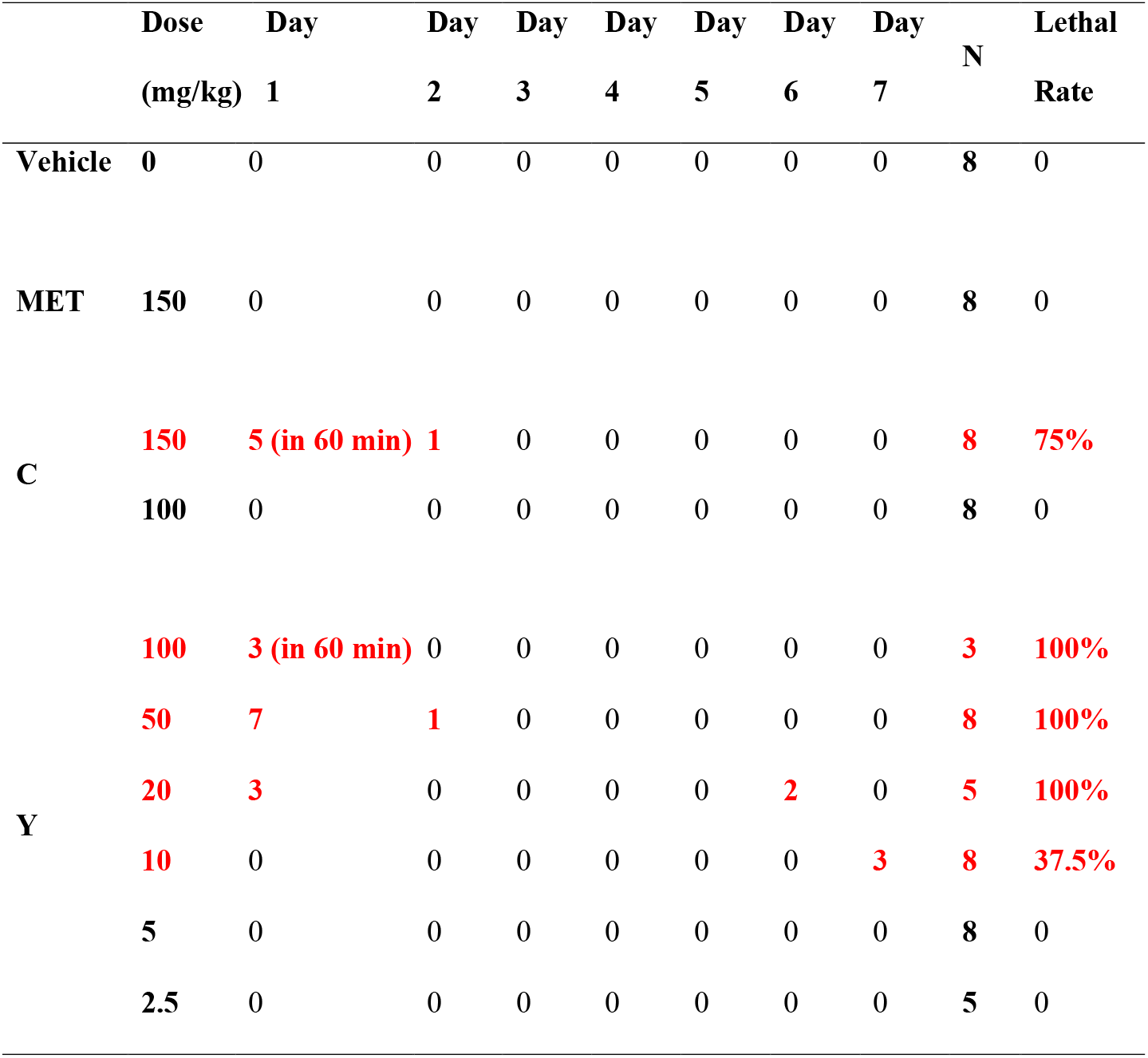
Lethality observations after single dose injection of compounds tested in one week. The treatment groups with death are highlighted in red. “N” is the number of mice used.

**Supplementary Table 2.**
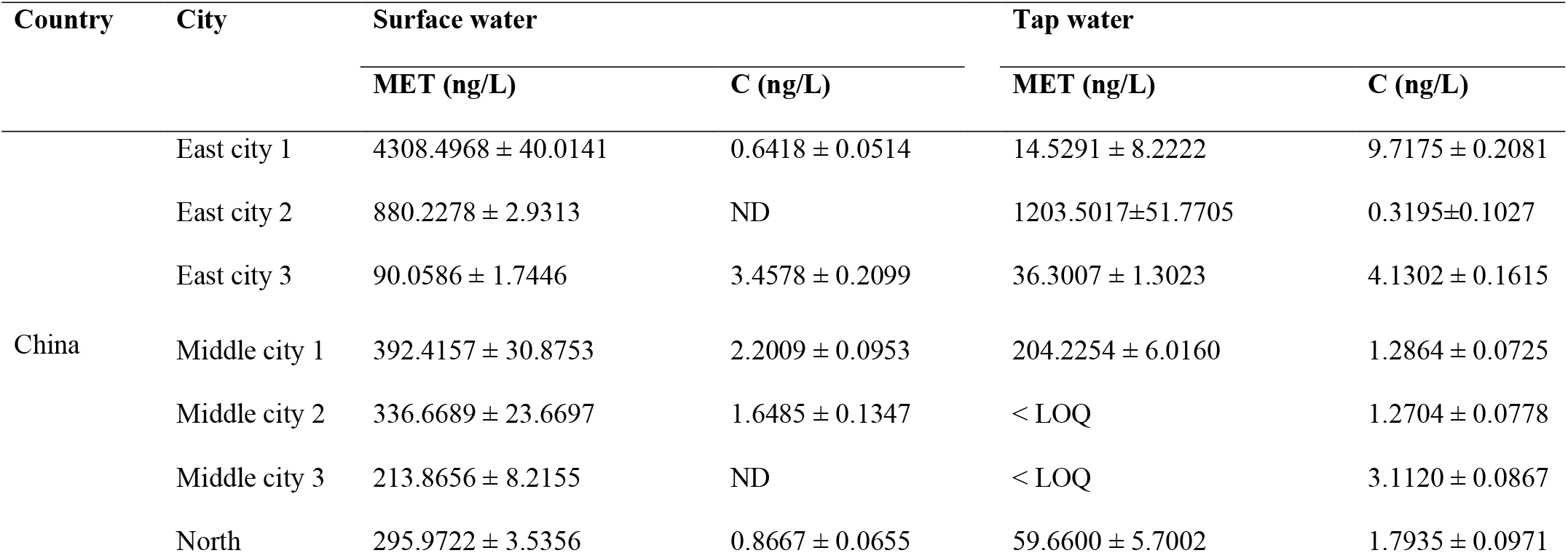
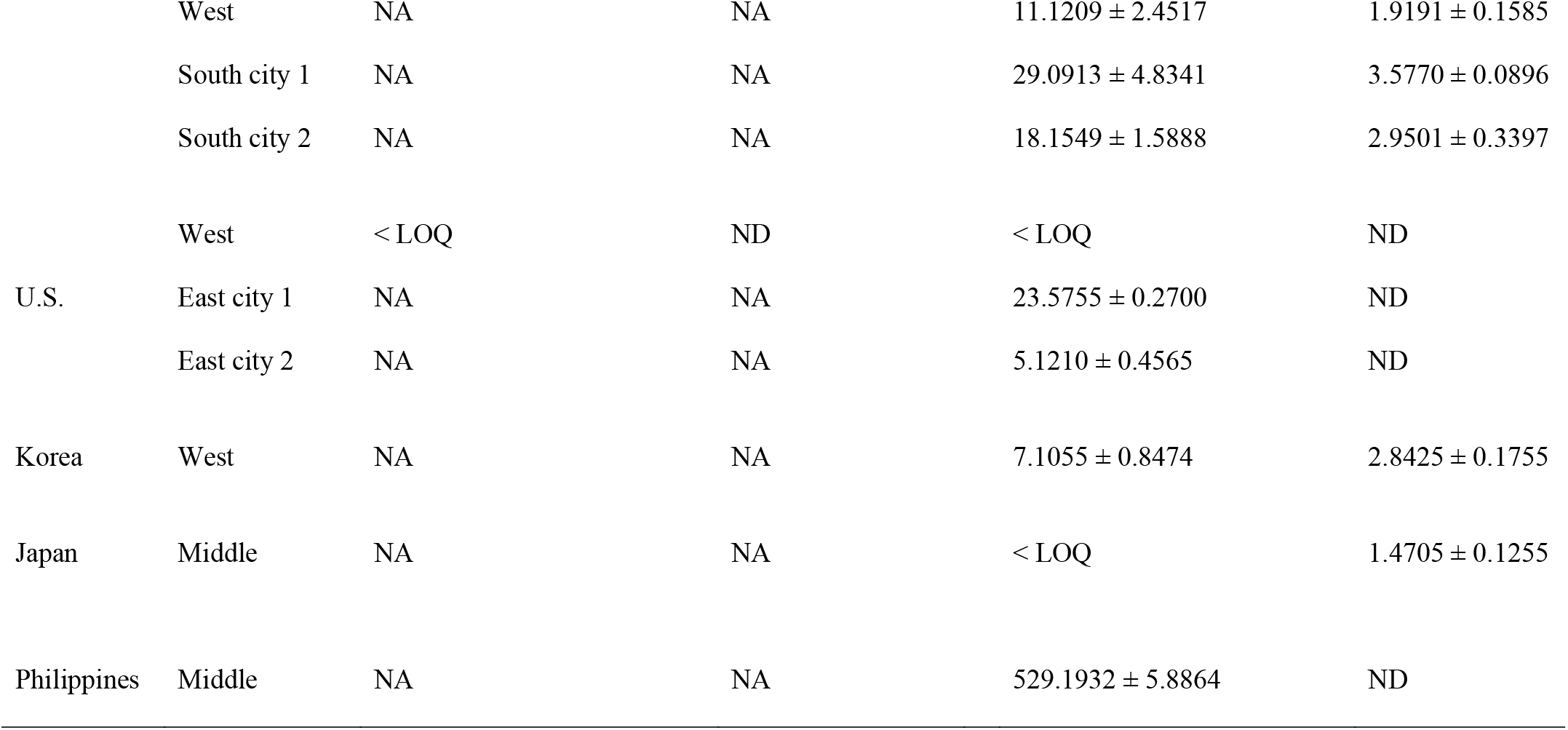
Current presences of MET and its byproduct C in surface water and tap water. We mainly sampled the reachable surface water, which serves as a drinking water source, and the tap water in different cities from several countries to investigate the concentrations of MET and byproducts Y and C. The data are presented as the mean value ± standard error of the mean (mean ± SEM) from three tests. The limit of quantitation (LOQ) of MET, byproduct Y and C are 5.00 ng/L, 90.75 ng/L and 1.00 ng/L, respectively. ND and <LOQ represent compounds that cannot be detected and are lower than the LOQ, respectively. NA means not available.

**Supplementary Table 3.**
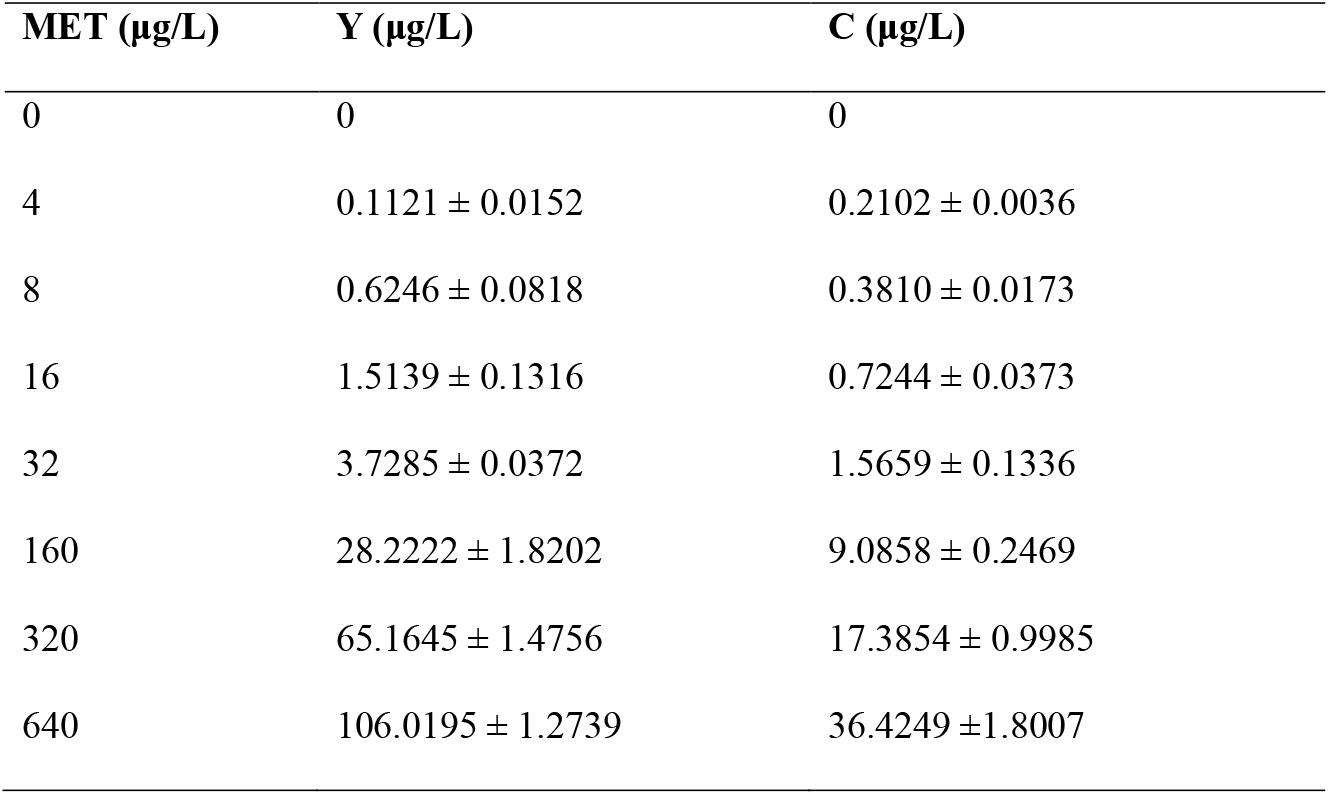
Simulated disinfection reaction at a fixed concentration of chlorine (0.3%) with increasing doses of MET. The first column shows the metformin concentrations used in the reaction. The yields of byproducts Y and C after 2 hours of reaction are provided as the mean value with standard error of the mean (mean ± SEM).

## References

1. W. C. Knowler, E. Barrett-Connor, S. E. Fowler, R. F. Hamman, J. M. Lachin, E. A. Walker, D. M. Nathan, G. Diabetes Prevention Program Research, Reduction in the incidence of type 2 diabetes with lifestyle intervention or metformin. N Engl J Med 346, 393–403 (2002).

2. A. a. o. Practice Committee of the American Society for Reproductive Medicine. Electronic address, M. Practice Committee of the American Society for Reproductive, Role of metformin for ovulation induction in infertile patients with polycystic ovary syndrome (PCOS): a guideline. Fertil Steril 108, 426–441 (2017).

3. A. A. Soukas, H. Hao, L. Wu, Metformin as Anti-Aging Therapy: Is It for Everyone? Trends Endocrinol Metab 30, 745–755 (2019).

4. L. Wu, B. Zhou, N. Oshiro-Rapley, M. Li, J. A. Paulo, C. M. Webster, F. Mou, M. C. Kacergis, M. E. Talkowski, C. E. Carr, S. P. Gygi, B. Zheng, A. A. Soukas, An Ancient, Unified Mechanism for Metformin Growth Inhibition in C. elegans and Cancer. Cell 167, 1705–1718 e1713 (2016).

5. ClinCalc. (ClinCalc DrugStats Database, 2020).

6. J. H. Yan, Y. Xiao, D. Q. Tan, X. T. Shao, Z. Wang, D. G. Wang, Wastewater analysis reveals spatial pattern in consumption of anti-diabetes drug metformin in China. Chemosphere 222, 688–695 (2019).

7. G. Rena, D. G. Hardie, E. R. Pearson, The mechanisms of action of metformin. Diabetologia 60, 1577–1585 (2017).

8. D. J. Caldwell, V. D’Aco, T. Davidson, K. Kappler, R. J. Murray-Smith, S. F. Owen, P. F. Robinson, B. Simon-Hettich, J. O. Straub, J. Tell, Environmental risk assessment of metformin and its transformation product guanylurea: II. Occurrence in surface waters of Europe and the United States and derivation of predicted no-effect concentrations. Chemosphere 216, 855–865 (2019).

9. P. L. Lena Stütz, Wolfgang Schulz and Rudi Winzenbacher Identification of genotoxic transformation products by effect-directed analysis with high-performance thin-layer chromatography and non-target screening. J Plan Chrom 32, 173–182 (2019).

10. R. D. MacLaren, K. Wisniewski, C. MacLaren, Environmental concentrations of metformin exposure affect aggressive behavior in the Siamese fighting fish, Betta splendens. PLoS One 13, e0197259 (2018).

11. E. Ussery, K. N. Bridges, Z. Pandelides, A. E. Kirkwood, J. Guchardi, D. Holdway, Developmental and Full-Life Cycle Exposures to Guanylurea and Guanylurea-Metformin Mixtures Results in Adverse Effects on Japanese Medaka (Oryzias latipes). Environ Toxicol Chem 38, 1023–1028 (2019).

12. R. M. Briones, W. Q. Zhuang, A. K. Sarmah, Biodegradation of metformin and guanylurea by aerobic cultures enriched from sludge. Environ Pollut 243, 255–262 (2018).

13. D. Armbruster, O. Happel, M. Scheurer, K. Harms, T. C. Schmidt, H. J. Brauch, Emerging nitrogenous disinfection byproducts: Transformation of the antidiabetic drug metformin during chlorine disinfection of water. Water Res 79, 104–118 (2015).

14. D. S. Lantagne, T. F. Clasen, Use of household water treatment and safe storage methods in acute emergency response: case study results from Nepal, Indonesia, Kenya, and Haiti. Environ Sci Technol 46, 11352–11360 (2012).

15. B. D. Blair, J. P. Crago, C. J. Hedman, R. D. Klaper, Pharmaceuticals and personal care products found in the Great Lakes above concentrations of environmental concern. Chemosphere 93, 2116–2123 (2013).

16. S. M. Elliott, M. E. Brigham, R. L. Kiesling, H. L. Schoenfuss, Z. G. Jorgenson, Environmentally relevant chemical mixtures of concern in waters of United States tributaries to the Great Lakes. Integr Environ Assess Manag 14, 509–518 (2018).

17. J. C. Florez, The pharmacogenetics of metformin. Diabetologia 60, 1648–1655 (2017).

18. S. A. Schmidt, E. Gukelberger, M. Hermann, F. Fiedler, B. Grossmann, J. Hoinkis, A. Ghosh, D. Chatterjee, J. Bundschuh, Pilot study on arsenic removal from groundwater using a small-scale reverse osmosis system-Towards sustainable drinking water production. J Hazard Mater 318, 671–678 (2016).

19. R. Mittal, C. M. Coopersmith, Redefining the gut as the motor of critical illness. Trends Mol Med 20, 214–223 (2014).

20. A. Kalender, A. Selvaraj, S. Y. Kim, P. Gulati, S. Brule, B. Viollet, B. E. Kemp, N. Bardeesy, P. Dennis, J. J. Schlager, A. Marette, S. C. Kozma, G. Thomas, Metformin, independent of AMPK, inhibits mTORC1 in a rag GTPase-dependent manner. Cell Metab 11, 390–401 (2010).

21. F. Cabreiro, C. Au, K. Y. Leung, N. Vergara-Irigaray, H. M. Cocheme, T. Noori, D. Weinkove, E. Schuster, N. D. Greene, D. Gems, Metformin retards aging in C. elegans by altering microbial folate and methionine metabolism. Cell 153, 228–239 (2013).

22. T. Ludman, O. K. Melemedjian, Bortezomib and metformin opposingly regulate the expression of hypoxia-inducible factor alpha and the consequent development of chemotherapy-induced painful peripheral neuropathy. Mol Pain 15, 1744806919850043 (2019).

23. M. Zang, A. Zuccollo, X. Hou, D. Nagata, K. Walsh, H. Herscovitz, P. Brecher, N. B. Ruderman, R. A. Cohen, AMP-activated protein kinase is required for the lipid-lowering effect of metformin in insulin-resistant human HepG2 cells. J Biol Chem 279, 47898–47905 (2004).

24. H. H. Gerets, K. Tilmant, B. Gerin, H. Chanteux, B. O. Depelchin, S. Dhalluin, F. A. Atienzar, Characterization of primary human hepatocytes, HepG2 cells, and HepaRG cells at the mRNA level and CYP activity in response to inducers and their predictivity for the detection of human hepatotoxins. Cell Biol Toxicol 28, 69–87 (2012).

25. P. R. Kiela, F. K. Ghishan, Physiology of Intestinal Absorption and Secretion. Best Pract Res Clin Gastroenterol 30, 145–159 (2016).

26. G. Hua, C. Wang, Y. Pan, Z. Zeng, S. G. Lee, M. L. Martin, A. Haimovitz-Friedman, Z. Fuks, P. B. Paty, R. Kolesnick, Distinct Levels of Radioresistance in Lgr5(+) Colonic Epithelial Stem Cells versus Lgr5(+) Small Intestinal Stem Cells. Cancer Res 77, 2124–2133 (2017).

27. H. Clevers, The intestinal crypt, a prototype stem cell compartment. Cell 154, 274–284 (2013).

28. T. Scholzen, J. Gerdes, The Ki-67 protein: from the known and the unknown. J Cell Physiol 182, 311–322 (2000).

29. A. H. Smith, P. A. Lopipero, M. N. Bates, C. M. Steinmaus, Public health. Arsenic epidemiology and drinking water standards. Science 296, 2145–2146 (2002).

30. L. Kong, K. Kadokami, S. Wang, H. T. Duong, H. T. C. Chau, Monitoring of 1300 organic micro-pollutants in surface waters from Tianjin, North China. Chemosphere 122, 125–130 (2015).

31. S. R. de Solla, E. A. Gilroy, J. S. Klinck, L. E. King, R. McInnis, J. Struger, S. M. Backus, P. L. Gillis, Bioaccumulation of pharmaceuticals and personal care products in the unionid mussel Lasmigona costata in a river receiving wastewater effluent. Chemosphere 146, 486–496 (2016).

32. M. Scheurer, A. Michel, H. J. Brauch, W. Ruck, F. Sacher, Occurrence and fate of the antidiabetic drug metformin and its metabolite guanylurea in the environment and during drinking water treatment. Water Res 46, 4790–4802 (2012).

33. C. A. Chiu, P. Westerhoff, A. Ghosh, GAC removal of organic nitrogen and other DBP precursors. J Am Water Works Ass 104, 41–42 (2012).

34. A. A. Cuthbertson, S. Y. Kimura, H. K. Liberatore, R. S. Summers, D. R. U. Knappe, B. D. Stanford, J. C. Maness, R. E. Mulhern, M. Selbes, S. D. Richardson, Does Granular Activated Carbon with Chlorination Produce Safer Drinking Water? From Disinfection Byproducts and Total Organic Halogen to Calculated Toxicity. Environ Sci Technol 53, 5987–5999 (2019).

35. A. H. Smith, E. O. Lingas, M. Rahman, Contamination of drinking-water by arsenic in Bangladesh: a public health emergency. Bull World Health Organ 78, 1093–1103 (2000).

36. M. Diana, M. Felipe-Sotelo, T. Bond, Disinfection byproducts potentially responsible for the association between chlorinated drinking water and bladder cancer: A review. Water Res 162, 492–504 (2019).

37. L. J. McCreight, C. J. Bailey, E. R. Pearson, Metformin and the gastrointestinal tract. Diabetologia 59, 426–435 (2016).

38. C. J. Bailey, C. Wilcock, J. H. Scarpello, Metformin and the intestine. Diabetologia 51, 1552–1553 (2008).

39. B. A. J. Poursat, R. J. M. van Spanning, M. Braster, R. Helmus, P. de Voogt, J. R. Parsons, Biodegradation of metformin and its transformation product, guanylurea, by natural and exposed microbial communities. Ecotoxicol Environ Saf 182, 109414 (2019).

40. F. Ju, K. Beck, X. Yin, A. Maccagnan, C. S. McArdell, H. P. Singer, D. R. Johnson, T. Zhang, H. Burgmann, Wastewater treatment plant resistomes are shaped by bacterial composition, genetic exchange, and upregulated expression in the effluent microbiomes. ISME J 13, 346–360 (2019).

## SI References

1. S. M. Elliott, M. E. Brigham, R. L. Kiesling, H. L. Schoenfuss, Z. G. Jorgenson, Environmentally relevant chemical mixtures of concern in waters of United States tributaries to the Great Lakes. Integr Environ Assess Manag 14, 509-518 (2018).

2. L. Kong, K. Kadokami, S. Wang, H. T. Duong, H. T. C. Chau, Monitoring of 1300 organic micro-pollutants in surface waters from Tianjin, North China. Chemosphere 122, 125–130 (2015).

3. S. R. de Solla, E. A. Gilroy, J. S. Klinck, L. E. King, R. McInnis, J. Struger, S. M. Backus, P. L. Gillis, Bioaccumulation of pharmaceuticals and personal care products in the unionid mussel Lasmigona costata in a river receiving wastewater effluent. Chemosphere 146, 486–496 (2016).

4. M. Scheurer, A. Michel, H. J. Brauch, W. Ruck, F. Sacher, Occurrence and fate of the antidiabetic drug metformin and its metabolite guanylurea in the environment and during drinking water treatment. Water Res 46, 4790–4802 (2012).

5. H. T. C. Chau, K. Kadokami, H. T. Duong, L. Kong, T. T. Nguyen, T. Q. Nguyen, Y. Ito, Occurrence of 1153 organic micropollutants in the aquatic environment of Vietnam. Environ Sci Pollut Res Int 25, 7147–7156 (2018).

6. T. L. ter Laak, P. J. Kooij, H. Tolkamp, J. Hofman, Different compositions of pharmaceuticals in Dutch and Belgian rivers explained by consumption patterns and treatment efficiency. Environ Sci Pollut Res Int 21, 12843–12855 (2014).

7. A. M. Ali, H. T. Ronning, W. Alarif, R. Kallenborn, S. S. Al-Lihaibi, Occurrence of pharmaceuticals and personal care products in effluent-dominated Saudi Arabian coastal waters of the Red Sea. Chemosphere 175, 505–513 (2017).

8. J. L. Schaper, M. Posselt, J. L. McCallum, E. W. Banks, A. Hoehne, K. Meinikmann, M. A. Shanafield, O. Batelaan, J. Lewandowski, Hyporheic Exchange Controls Fate of Trace Organic Compounds in an Urban Stream. Environ Sci Technol 52, 12285–12294 (2018).

9. E. E. Burns, L. J. Carter, D. W. Kolpin, J. Thomas-Oates, A. B. A. Boxall, Temporal and spatial variation in pharmaceutical concentrations in an urban river system. Water Res 137, 72–85 (2018).

10. N. A. Al-Odaini, M. P. Zakaria, M. I. Yaziz, S. Surif, M. Abdulghani, The occurrence of human pharmaceuticals in wastewater effluents and surface water of Langat River and its tributaries, Malaysia. Int J Environ an Ch 93, 245–264 (2013).

11. N. Park, Y. Choi, D. Kim, K. Kim, J. Jeon, Prioritization of highly exposable pharmaceuticals via a suspect/non-target screening approach: A case study for Yeongsan River, Korea. Sci Total Environ 639, 570–579 (2018).

12. A. S. Kachhawaha, P. M. Nagarnaik, M. Jadhav, A. Pudale, P. K. Labhasetwar, K. Banerjee, Optimization of a Modified QuEChERS Method for Multiresidue Analysis of Pharmaceuticals and Personal Care Products in Sewage and Surface Water by LC-MS/MS. J AOAC Int 100, 592–597 (2017).

13. M. Ruff, M. S. Mueller, M. Loos, H. P. Singer, Quantitative target and systematic non-target analysis of polar organic micro-pollutants along the river Rhine using high-resolution mass-spectrometry--Identification of unknown sources and compounds. Water Res 87, 145–154 (2015).

14. E. Vulliet, C. Cren-Olive, Screening of pharmaceuticals and hormones at the regional scale, in surface and groundwaters intended to human consumption. Environ Pollut 159, 2929–2934 (2011).

15. T. P. Wood, C. Du Preez, A. Steenkamp, C. Duvenage, E. R. Rohwer, Database-driven screening of South African surface water and the targeted detection of pharmaceuticals using liquid chromatography - High resolution mass spectrometry. Environ Pollut 230, 453–462 (2017).

16. P. Xu, J. E. Drewes, C. Bellona, G. Amy, T. U. Kim, M. Adam, T. Heberer, Rejection of emerging organic micropollutants in nanofiltration-reverse osmosis membrane applications. Water Environ Res 77, 40–48 (2005).

17. C. Lindim, J. van Gils, D. Georgieva, O. Mekenyan, I. T. Cousins, Evaluation of human pharmaceutical emissions and concentrations in Swedish river basins. Sci Total Environ 572, 508–519 (2016).

18. https://diabetesatlas.org/en/resources/. (2019).

